# Negative selection in humans and fruit flies involves synergistic epistasis

**DOI:** 10.1101/066407

**Authors:** Mashaal Sohail, Olga A. Vakhrusheva, Jae Hoon Sul, Sara Pulit, Laurent Francioli, GoNL Consortium, Alzheimer’s Disease Neuroimaging Initiative, Leonard H. van den Berg, Jan H. Veldink, Paul de Bakker, Georgii A. Bazykin, Alexey S. Kondrashov, Shamil R. Sunyaev

**Affiliations:** Systems Biology PhD Program, Harvard Medical School, Boston, MA, USA; Division of Genetics, Department of Medicine, Brigham and Women’s Hospital, Harvard Medical School, Boston, MA, USA; Institute for Information Transmission Problems (Kharkevich Institute) of the Russian Academy of Sciences, Moscow, Russia; Pirogov Russian National Research Medical University, Moscow, Russia; Department of Psychiatry and Biobehavioral Sciences, UCLA, Los Angeles, CA, USA; Department of Neurology and Neurosurgery, Brain Center Rudolf Magnus, University Medical Center Utrecht, Utrecht, the Netherlands; Department of Medical Genetics, Center for Molecular Medicine, University Medical Center Utrecht, Utrecht, the Netherlands; Skolkovo Institute of Science and Technology, Skolkovo, Russia; Department of Bioengineering and Bioinformatics, M.V. Lomonosov Moscow State University, Moscow, Russia; Department of Ecology and Evolutionary Biology, University of Michigan, Ann Arbor, MI, USA; Broad Institute of Harvard and MIT, Cambridge, MA, USA

**Keywords:** Epistasis, negative selection, evolution, sex, genetic recombination, mutation load, linkage disequilibrium

## Abstract

Negative selection against deleterious alleles produced by mutation is the most common form of natural selection, which strongly influences within-population variation and interspecific divergence. However, some fundamental properties of negative selection remain obscure. In particular, it is still not known whether deleterious alleles affect fitness independently, so that cumulative fitness loss depends exponentially on the number of deleterious alleles, or synergistically, so that each additional deleterious allele results in a larger decrease in relative fitness. Negative selection with synergistic epistasis must produce negative linkage disequilibrium between deleterious alleles, and therefore, underdispersed distribution of the number of deleterious alleles in the genome. Indeed, we detected underdispersion of the number of rare loss-of-function (LoF) alleles in eight independent datasets from modern human and *Drosophila melanogaster* populations. Thus, ongoing selection against deleterious alleles is characterized by synergistic epistasis, which can explain how human and fly populations persist despite very high genomic deleterious mutation rates.

---

Negative selection plays a key role in evolution, preventing unlimited accumulation of deleterious mutations and establishing the mutation-selection equilibrium (1). The properties of negative selection are determined by the corresponding fitness landscape, the function which relates fitness to the “mutation burden” of a genotype. In the simplest case of equally deleterious mutations, mutational burden is the total number of mutant alleles in a genome. According to the null hypothesis of no epistasis, selection acts on different mutations independently, so that each additional mutation causes the same decline in relative fitness and fitness depends exponentially on their number. By contrast, if synergistic, or narrowing (2), epistasis between deleterious alleles is present, each additional mutation causes a larger decrement of relative fitness. Synergistic epistasis can reduce the mutation load under a given genomic rate of deleterious mutations (1, 3–4) and can produce the evolutionary advantage of sex and recombination (5). However, because neither the mutational burden nor fitness can be easily measured, data on fitness landscapes of negative selection remain inconclusive (6). Recent genome-wide investigations have found pervasive epistasis, but no consistent directionality of effect (6–8). Synergistic epistasis between deleterious mutations is more prevalent in organisms with complex genomes (7). Moreover, theoretical work suggests that narrowing epistasis may emerge as a result of pervasive pleiotropy, and modular organization of biological networks (9). This would lead to antagonism between beneficial mutations and synergism between deleterious mutations.

In this paper, we study the distribution of the mutation burden in human and *Drosophila melanogaster* populations. In the absence of epistasis, deleterious alleles independently contribute to the mutation burden (3). Thus, if mutant alleles are rare, the mutation burden has Poisson distribution, so that its variance (*σ*^2^) is equal to its mean (*μ*) (Fig. S1). More generally, the variance of the mutation burden is equal to the sum of variances at all deleterious loci, or the additive variance (*V_A_*) (10), computed as Σ_*i*_ 2*p_i_*(1 − *p_i_*) for all deleterious loci *i* with mutant allele frequency *p_i_* in the genome (Fig. 1). This is mathematically equivalent to the genome-wide nucleotide diversity of deleterious alleles (11). In contrast, epistatic selection creates dependencies between individual alleles, so total variance of the mutation burden is no longer equal to the additive variance. Selection with synergistic epistasis creates negative linkage disequilibria (LDs) between deleterious alleles. Due to the dependencies between individual alleles, variance of the mutation burden is reduced by a factor of *ρ* (<1), which is determined by the strength of selection and the extent of epistasis (13, 14, Fig. S2). Antagonistic epistasis, instead, creates positive linkage disequilibria (LDs) between deleterious alleles and increases variance of the mutation burden. Truncation selection, which represents the extreme mode of synergistic epistasis (4) leads to the smallest *ρ*. In the extreme example, if 50% of individuals with above average numbers of mutations would not contribute to the next generation, *ρ* = 0.36 if the average genomic number of mutations is high. Because free recombination halves LDs in the course of one generation, at the mutation-selection equilibrium *σ*^2^ = *V_A_*/2 − *ρ*, where *V_A_* is the variance of the mutation burden under linkage equilibrium. More subtly, the difference between *σ*^2^ and *V_A_* is also a genome-wide estimate of the net linkage disequilibrium in fitness. For all pairs of loci *i* and *j* in the genome, *I* = *σ*^2^ − *V_A_* = 4Σ*_i,j_D_i,j_*, where *D_i,j_* is the pair-wise linkage disequilibrium. Using data on multiple genotypes from a population, we utilized this statistical framework to create a test for synergistic epistasis without the need to measure fitness.

**Fig 1.**
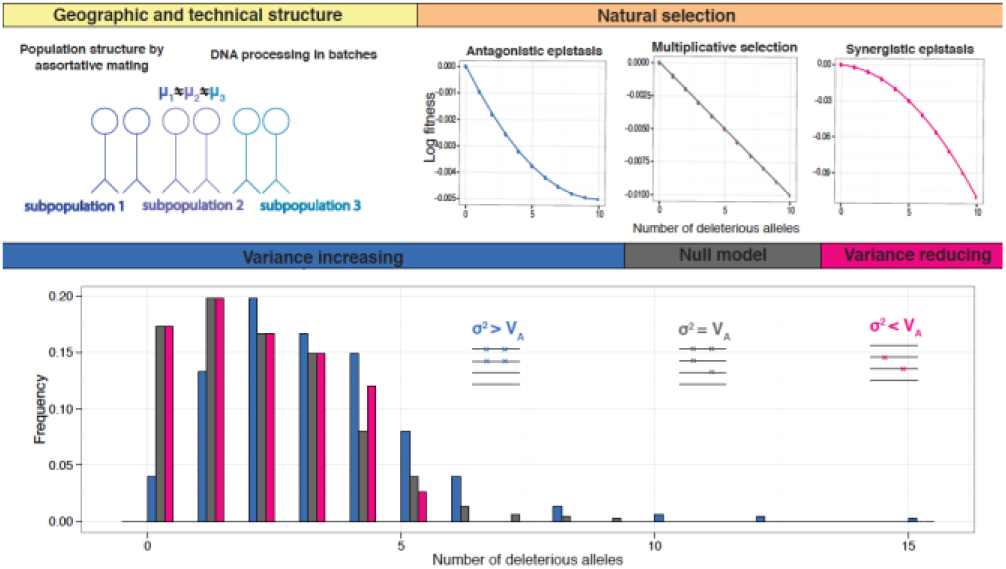
Rare mutation burden under natural selection (*orange, right*) and population structure (*yellow, left*). In the absence of epistasis (grey), variance (*σ*^2^) of the mutation burden (bottom panel) is equal to its additive variance (*V_A_*). Overdispersion (*blue*) occurs due to natural selection with antagonistic epistasis, and due to population structure and technical heterogeneity during sequencing. Underdispersion (*pink*) occurs due to natural selection with synergistic epistasis.

The ideal population for our test would be single ancestry, out-breeding, non-admixed and randomly mating. We analyzed three suitable European datasets – the Genome of the Netherlands (GoNL) (14), Alzheimer’s Disease Neuroimaging Initiative (ADNI), and Dutch controls from Project MinE, an amyotrophic lateral sclerosis (ALS) study. For each of these, we obtained whole-genome sequences of unrelated individuals. We obtained the same data for Zambian flies from Phase 3 of the *Drosophila* Population Genomics Project (DPGP3) (15). For each population, after applying stringent quality control filters (Tables S8-S12), we computed the mutation burden distribution for coding synonymous, missense, and loss-of-function or LoF, defined as splice site disrupting and nonsense variants. For all of these datasets, distribution of LoF singletons was underdispersed (nonsense variants in the MinE dataset, if considered separately, were the exception, although underdispersion was also observed for stop gain variants in this dataset at a slightly higher allele frequency threshold (Table S2)). On average, rare LoF variants displayed variance (*σ*^2^) reduced by a factor of ~0.9, or underdispersion, compared to additive variance (*V_A_*) (Table 1, Fig. 2). Thus, *σ*^2^ = 0.9*V_A_* is consistent with *ρ* = 0.89 which appears, for example, after truncation of less than 2% of the population. In contrast, rare coding synonymous variants showed *σ*^2^ greater than *V_A_*, or overdispersion. We replicated the same signal in three non-European populations from the 1000 genomes Phase I Project (16) (Table S1, Table S2) and an American *D. melanogaster* population from the *D. melanogaster* Genetic Reference Panel (DGRP, *17*, Table S3). We proceeded to ask two questions: why were the synonymous variants overdispersed compared to their expectation under independence? Was the underdispersion in LoF variants significant?

**Fig 2.**
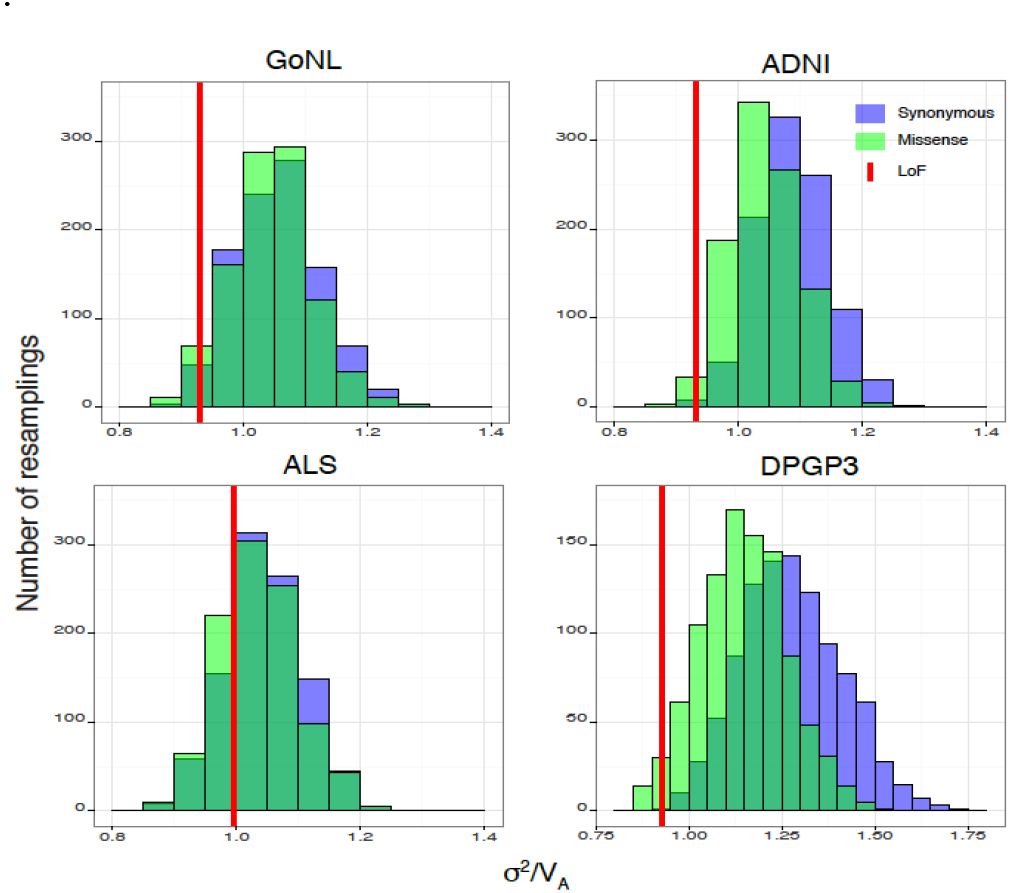
Resampling distributions of *σ*^2^ to *V_A_* ratio (*n*) for LoF rare mutation burden in humans and *D. melanogaster*. Synonymous (purple) and missense (green) variants were resampled at the same mean and allele frequency as LoF variants to obtain empirical null distributions for the variance *σ*^2^ to additive variance *V_A_* ratio for each dataset. For humans, only singletons are included (see Table S2 for other frequency cut-offs). For flies, alleles up to a minor allele count of 5 are included (see Table S3 for other cut-offs). A synonymous p-value for the *σ*^2^ to *V_A_* ratio of the rare LoF mutation burden (red) was obtained for each dataset (Table S1), and a joint p-value for all 3 human datasets shown (GoNL, ADNI, ALS) was computed by meta-analysis using Stouffer’s method (p = 0.0003).

**Table 1.**
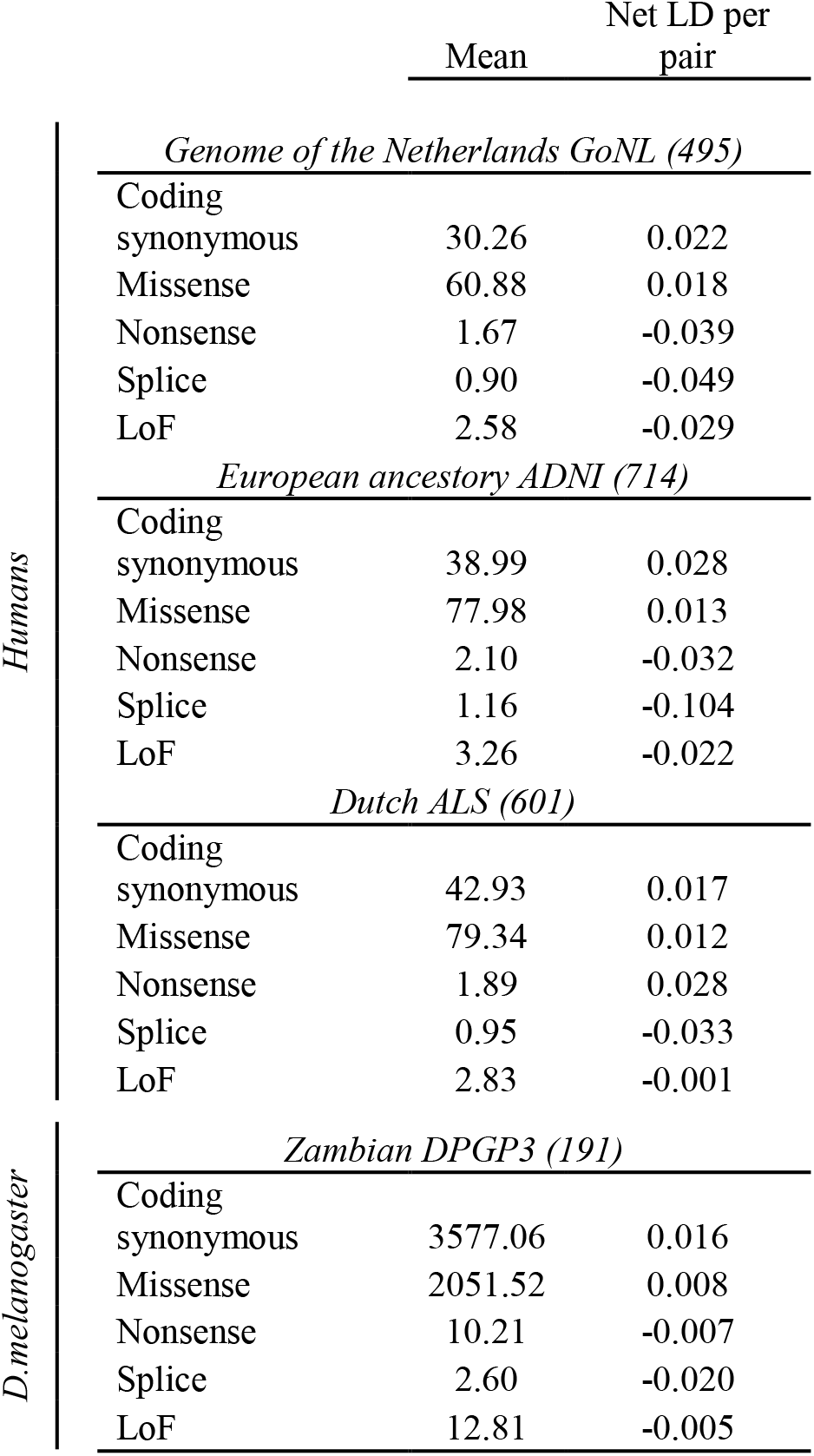
Negative linkage disequilibrium (LD) between rare LoF variants in human and *D. melanogaster* genomes. For humans, only singletons are included (see Table S2 for other frequency cut-offs). For flies, alleles up to a minor allele count of 5 are included (see Table S3 for other cut-offs). The number of samples is given in parentheses for each dataset. LoF variants include splice site disrupting and nonsense variants. Net LD per pair of alleles (*Í*) is computed as the difference between the variance *σ*^2^ and additive variance *V_A_* of the rare mutation burden, normalized by the square of the mean mutation burden μ (*Í* = *I*/*μ*^2^). A p-value was obtained for each human dataset by permutation (Table S1), and a joint p-value for all 3 human datasets shown (GoNL, ADNI, ALS) was computed by meta-analysis using Stouffer’s method (coding synonymous p=0.999, missense p < 1x10^−3^, LoF p = 0.002).

Even for a set of independent alleles, overdispersion in the mutation burden is observed if genome-wide positive LD is present due to population structure, which can also be seen as deviations from Hardy-Weinberg equilibrium for the entire genome (Fig. 1, 18, 19). If the population has a cline in average mutation burden (*μ*)(20) due to, for example, a south-to-north expansion (14)(21) followed by assortative mating, this translates into an excess of *σ*^2^ over *V_A_* (Fig. S3, Fig. S4, Table S4). Overdispersion can also be caused by technical reasons (Fig. 1). When DNA samples are sequenced or processed in different batches, the heterogeneity introduced can result in a clustering effect similar to that of geographic structure. Using GoNL samples, for which we had detailed geographic and technical information, we showed that a large proportion of the overdispersion in rare mutation burden could be attributed to geographic origin and sequencing batch (Fig. S5, Table S4). We also showed that by calculating mutation burden across allele frequencies, where no strong differences in *μ* between human populations have been detected (20), we effectively observed an independent distribution of mutation burden for all variants (Table S2).

We next proceeded to investigate the contrast between coding synonymous and LoF variants. Having uncovered the primary sources for overdispersion of rare mutation burden, we realized that overdispersion scaled with the mean (*μ*) of the mutation burden distribution (Table S5). We, therefore, generated an empirical null distribution for each dataset by resampling coding synonymous variants at the same mean (*μ*) and allele frequency as our test set of LoF variants (1000 resamples for each dataset, Fig. 2). Meta-analyzing across all datasets using Stouffer’s method (Tables S1-S3), we showed that the deleterious mutation burden for LoF variants was significantly underdispersed in humans (p = 0.0003) and flies (p = 9.43 x 10^−6^). We also tested significance in humans using an alternative approach (Table S1). Permuting functional consequences across variants, we confirmed the significance of our underdispersion signal in deleterious mutation burden (missense p < 1x10^−3^, LoF p = 0.002). Furthermore, we showed that the underdispersion signal persists after correcting raw metrics for population structure and other confounding factors (Table S5)

Notably, the detected signal varies between human and fly populations. The underdispersion signal in deleterious mutation burden is stronger in flies compared to humans (Fig. 2). First, this is because recombination, which opposes the reduction in genetic variance caused by negative LDs, is weaker in flies compared to humans. The observed effect decreases with the harmonic mean cH of the recombination frequencies among the sites involved (22). Flies, with only four pairs of chromosomes and no crossing over in males, has an estimated cH of 0.1, significantly lower than a cH close to 0.4 for humans (23). Second, in industrialized human populations, after the second demographic transition, selection due to pre-reproductive mortality is deeply relaxed (24, 25). Thus, recombination would rapidly destroy linkage disequilibria between deleterious alleles.

While several factors – geographic structure, technical issues, antagonistic epistasis – can lead to an overdispersed mutation burden distribution, only synergistic epistasis can lead to an underdispersed distribution for unlinked loci. We further partitioned the underdispersion signal into within- and between-chromosome components, and demonstrated that the deleterious mutation burden was underdispersed due to multilocus associations both within and between different chromosomes (Table S6, Table S7). While the negative linkage disequilibria from linked regions may be attributed to Hill-Robertson interference(26, 27) the majority of our underdispersion signal comes from unlinked pairs of loci, even for pairs of loci within the same chromosome (Fig. S6, Fig. S7). Having identified and controlled for other sources of LD, we thus invoke synergistic epistasis as a significant contributor to the underdispersion signal in deleterious mutation burden that we observe in humans and flies.

Thirty years ago, Neel posed the question: “The amount of silent DNA is steadily shrinking. The question of how our species accommodates such [high deleterious] mutation rates is central to evolutionary thought (28).” Indeed, a newborn receives ~70 *de novo* mutations (29). Although estimates for the target size for deleterious alleles (fraction of the genome that is “functional”) vary, an overwhelming majority suggest that about 10% of the human genome sequence is functionally significant and selectively constrained (30, 31). Thus, the average human individual is expected to carry at least seven *de novo* deleterious mutations, which is incompatible with the long-term population survival if selection is non-epistatic. Moreover, regardless of epistasis, at the mutation-selection equilibrium, the sum of coefficients of selection against mutant alleles present in an average genotype must equal the genome-wide deleterious mutation rate (32). Recently Henn et al (20) independently estimated this sum in humans to be 15. Without epistatic selection, this suggests a mutation load that is inconsistent with the existence of the population (1-e^−15^ >0.999). Thus, synergistic epistasis is the only way for humans to survive, and in a sense, our findings are not unexpected. In industrialized human populations, while selection due to pre-reproductive mortality is deeply relaxed, there is still a substantial opportunity for selection due to differential fertility (33). Also, only ~30% of human conceptions result in live births (34), indicating a substantial opportunity for prenatal selection. Thus, our results suggest that epistatic negative selection in humans is ongoing.

## Acknowledgements

We would like to thank Leonid Mirny, Gill McVean, Rong-Cai Yang and Ivan Adzhubey for useful scientific discussions. We would like to thank Onuralp Soylemez, Winston Anthony, David Radke and members of Sunyaev lab for comments on the manuscript. The authors are grateful to Dr. Justin Lack for help with *D. melanogaster* inversion data.

The project was supported by NIH grants R01GM078598, R01GM105857, R01MH101244. Analysis of fruit fly data was performed at IITP RAS and supported by the Russian Science Foundation grant no. 14-50-00150.

Part of the data collection and sharing for this project was funded by the Alzheimer’s Disease Neuroimaging Initiative (ADNI) (National Institutes of Health Grant U01 AG024904) and DOD ADNI (Department of Defense award number W81XWH-12-2-0012). ADNI is funded by the National Institute on Aging, the National Institute of Biomedical Imaging and Bioengineering, and through generous contributions from the following: AbbVie, Alzheimer’s Association; Alzheimer’s Drug Discovery Foundation; Araclon Biotech; BioClinica, Inc.; Biogen; Bristol-Myers Squibb Company; CereSpir, Inc.; Cogstate; Eisai Inc.; Elan Pharmaceuticals, Inc.; Eli Lilly and Company; EuroImmun; F. Hoffmann-La Roche Ltd and its affiliated company Genentech, Inc.; Fujirebio; GE Healthcare; IXICO Ltd.; Janssen Alzheimer Immunotherapy Research & Development, LLC.; Johnson & Johnson Pharmaceutical Research & Development LLC.; Lumosity; Lundbeck; Merck & Co., Inc.; Meso Scale Diagnostics, LLC.; NeuroRx Research; Neurotrack Technologies; Novartis Pharmaceuticals Corporation; Pfizer Inc.; Piramal Imaging; Servier; Takeda Pharmaceutical Company; and Transition Therapeutics. The Canadian Institutes of Health Research is providing funds to support ADNI clinical sites in Canada. Private sector contributions are facilitated by the Foundation for the National Institutes of Health (www.fnih.org). The grantee organization is the Northern California Institute for Research and Education, and the study is coordinated by the Alzheimer’s Therapeutic Research Institute at the University of Southern California. ADNI data are disseminated by the Laboratory for Neuro Imaging at the University of Southern California.

This study/database makes use of data generated by the Genome of the Netherlands Project. A full list of the investigators is available from http://www.nlgenome.nl. Funding for the project was provided by the Netherlands Organization for Scientific Research under award number 184021007, dated July 9, 2009 and made available as a Rainbow Project of the Biobanking and Biomolecular Research Infrastructure Netherlands (BBMRI-NL). The sequencing was carried out in collaboration with the Beijing Institute for Genomics (BGI).

